# Serial Two-Photon Tomography Imaging of the whole marmoset brain for neuroanatomical analyses

**DOI:** 10.1101/2024.09.03.611138

**Authors:** Akiya Watakabe, Toshiki Tani, Hiroshi Abe, Henrik Skibbe, Noritaka Ichinohe, Tetsuo Yamamori

## Abstract

Serial Two photon tomography (STPT) imaging is a technique to image a mass of tissue in its three-dimensional shape by combining two-photon imaging with automatic stage control and microtome slicing. We successfully implemented this technique for tracing the axonal projections in the marmoset brain. Here we describe the detailed experimental procedures that realized reliable volumetric imaging of the whole marmoset brain. We also introduce auxiliary histological techniques including visualization of myelin structure by simple light reflection, and immunological detection of non-fluorescent anterograde and retrograde tracers, which further enhance the utility of STPT data.

## INTRODUCTION

In neuroanatomical studies, researchers are faced with the need to observe μm-order structures (e.g., axons and boutons) in the context of the whole brain. This difficult task has been typically approached by visually inspecting the serial sections and looking for the region of interest for detailed imaging, analyses, and photo recording. With technological advancement, however, it’s becoming possible to image the entire brain at high resolution for whole-brain analysis. In a ground-breaking study by Oh et al.^1^, hundreds of mouse brains received tracer injections for connectomic analysis, which were processed by Serial Two-photon Tomography (STPT) imaging technique^2^. What was quite characteristic of this study was that they selected anterograde tracers for the quantification of “connectivity”. While anterograde tracers provide very detailed spatial information on axonal distribution, neuroanatomists have relied on laborious manual segmentation for its analysis. By automating various analytical procedures including this segmentation step, they succeeded in the “mass production” of ready-to-use high-resolution tracer data for multi-purpose uses. The effectiveness of their approach is obvious given a variety of studies that used their tracer data^3–5^ and their standard brain ^6^.

Historically, the neural connectivity of non-human primate brains has attracted the attention of many neuroanatomists since the development of the degeneration method in the 1950s, to anterograde/retrograde substance transportation methods in the 1970s to the present viral strategy^7, 8^. As such, there exist vast pieces of literature that investigated primate neural connections. In particular, many researchers investigated the complex corticocortical connectivity of macaque brain and their results have been curated for tabulation (e.g., CoCoMac^9^). Although useful, these classic studies had several limitations. First, because each study focuses only on limited brain region, the obtained information inevitably becomes fragmental. Second, each study uses different methods and conditions. Thus, the quantitative evaluation across studies become complicated. Third, the connectivity is usually shown either as camera lucida of representative sections or as semi-quantitative table/graph for author-defined brain regions. In other words, only a very limited information out of a complex brain structure is extracted for presentation in the published literatures. With the development of MRI techniques, the whole-brain studies at low resolution became prominent. However, there is a large gap between the levels of details between what we know about neural connections in mice and those in primates.

With such background in mind, we set out to perform comprehensive anterograde tracing of the common marmoset brains^10, 11^. Although much smaller and smoother than the macaque brain, the marmoset counterpart exhibits clear signs of primates, such as the presence of area MT and the granular prefrontal areas, neither of which is clearly defined in rodents^12, 13^. Here, the small size was a big advantage, because even the marmoset brain weighs 10 times greater than the mouse brain. Fortunately, we could image the whole marmoset brain with minimal software updates of TissueCyte (TissueVision Inc.), which is the commercially available microscope for STPT imaging^2^. In this article, we share our protocol to handle the marmoset brain for STPT imaging. We also provide the post-imaging protocol that further enhances the utility of this method.

## PROTOCOL

### 0. Tracer injection

1. Inject fluorescent and non-fluorescent tracers to the marmoset brain^14^. As the fluorescent tracer, we recommend using a double-vector TET-Off system^10^ for enhanced expression, because STPT requires detection of the native fluorescence. We routinely used Biotinylated Dextran Amine (BDA) as the non-fluorescent tracer. We also used cre-expressing AAV in AAV2 retro for retrograde detection of the input cell nuclei^10^. We also developed GFP-based smFP-tag AAVs as the non-fluorescent tracers (see below).
2. Perfusion fix and obtain the marmoset brain after four weeks.
3. Keep the brain in 4% paraformaldehyde/0.1M PB for 48 hours at 4C and transfer to 50 mM PB. If not used immediately, store the brain in 0.75% Glycine/0.1M PB to prevent autofluorescence due to overfixation.

### 1. Sample preparation

1. Incubate the fixed brain with collagenase (Wako chemical; collagenase type I ; #031-17601) at 37C for one hour (1mg/ml in 3mM CaCl2 in 10 ml TBS).
2. Carefully rub the brain surface with a cotton swab to peel off the pia matter (Fig. 1A). Use fine forceps to peal the detached meninges. It’s difficult to expose these structures that are hidden deep inside. But you need to remove meninges as much as possible. Otherwise, they often remain uncut, float up and interfere with imaging and slicing.
3. In our case, we routinely performed ex vivo MRI, which uses fluorine solution^15^. The leftover of the fluorine, if embedded together, could form tiny air bubbles under the objective while imaging. To avoid this, keep the brain in 0.1M phosphate buffer for one week before embedding. Otherwise, you can immediately proceed to embedding.
4. Prepare the NaBH4 buffer by dissolving 0.2 g NaBH4 (Sigma, Catalog # 452882) in 100 ml of 50 mM borate buffer (pH9.2) warmed to 40C. Leave the cap loose to let the CO2 gas out. Keep the solution overnight with the cap loosen.
5. Make oxidized agarose by stirring 2.25 g agarose (Sigma, Type 1, Catalog # A6013) and 0.21 g NaIO4 (Sigma, Catalog # S1878) in 100 ml phosphate buffer (PB) for 2-3 hours. Filter solution with vacuum suctioning and wash with three changes of the phosphate buffer (PB). Resuspend the agarose in 50 ml PB.
6. Completely melt the agarose by microwave oven and cool down to 60-65C.
7. Place the brain into a custom-made chamber and embed the brain in agarose. Take care to introduce the agarose into the cavity below the corpus callosum (Fig. 1B).
8. Disassemble the chamber and immerse the agarose block in the NaBH4 buffer overnight at 4C.
9. Change the buffer to PB several times over one to two weeks at 4C. The tissue autofluorescence becomes very weak if the buffer exchange is incomplete.
10. Make a slideglass stage by attaching four Neodymium Magnets with epoxy instant mix (Locktite).
11. Mount the agarose block onto the stage with superglue.

**Fig. 1:**
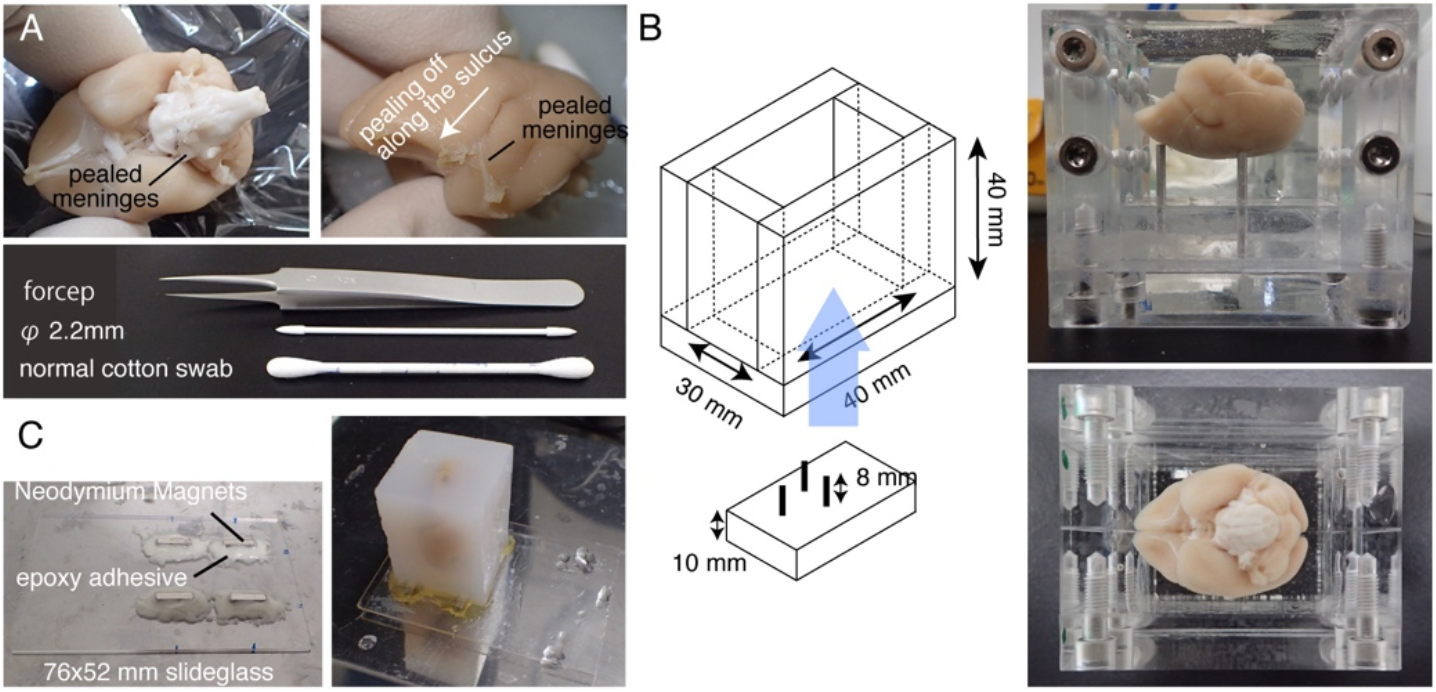
Sample preparation for STPT. (A) Meninge removal using cotton swabs. The meninges surrounding the brainstem can be removed by fine-tipped forceps. The meninges surrounding the marmoset brain is removed manually by rubbing with cotton swabs. The photo shows the meninges pealed from within the lateral sulcus. (B) The acryl box used for agarose embedding. The pins are movable and used to adjust the angle of the brain to be close to the stereotaxically fixed position^26^. (C) A magnetic slide made of 76×52 mm slideglass and four Neodymium magnets attached by epoxy adhesive. The agarose block is attached to the magnetic stage with superglue.

### 2. TissueCyte processing

1. Operate TissueCyte according to manufacturer’s instruction. A detailed protocol is also published^16^.
2. Several key points for treating marmoset brains are described below.

- The entire marmoset brain can be processed coronally from the anterior to the posterior ends without any changes in the hardware. But caution is needed not to exceed the stage moving limits when placing the brain.
- The angle of the blades needs to be finely tuned so that the surface depths at both ends of the blade is within 10 μm. Because of dense myelination, even 10 μm difference in depth could lead to different laser penetration through adult marmoset white matter. With the same reason, we usually set the imaging plane at about 25-35 μm from the surface.
- Ceramic blade is recommended to cut the entire brain at 50 μm interval (∼over 650 slices).
- The meninges remain uncut and interfere with slicing if not properly removed (Fig. 2). Some meninges (e.g., those surrounding the hippocampus or pulvinar) are difficult to remove.

**Fig. 2:**
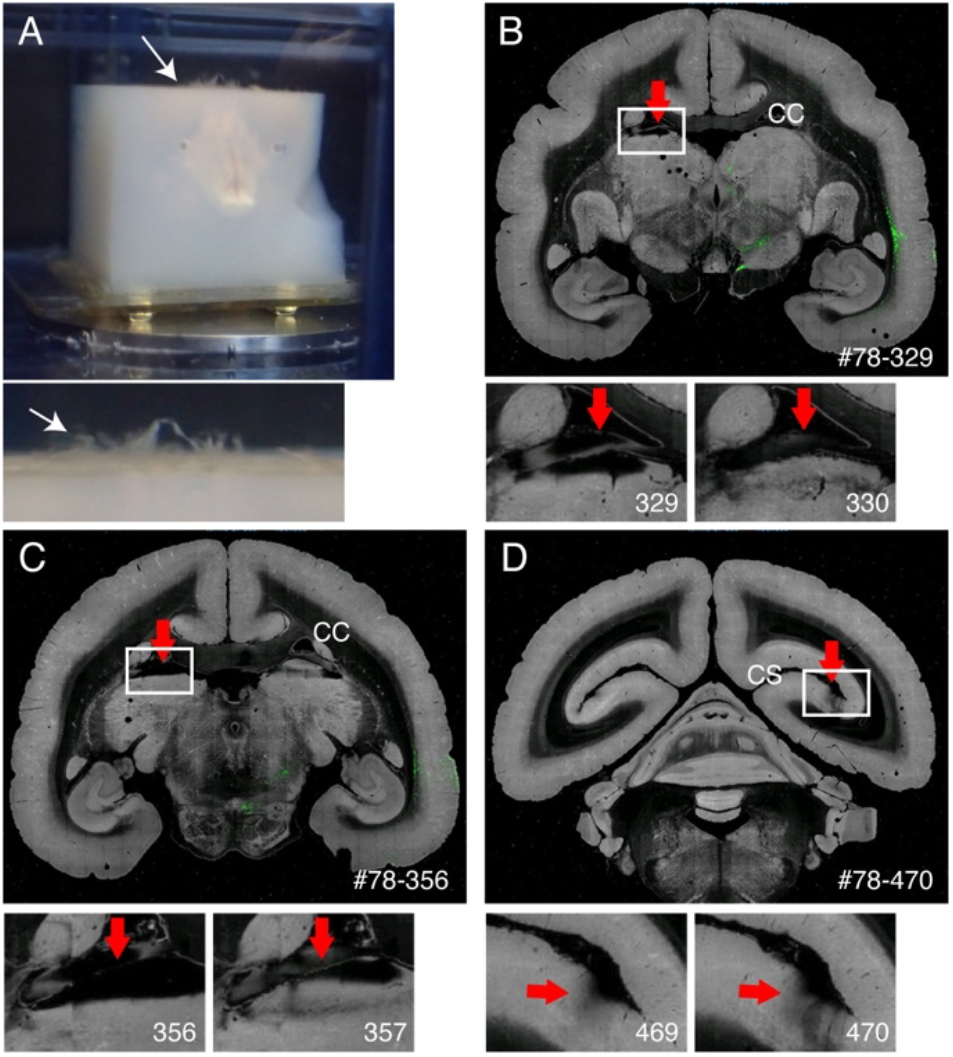
The effect of meninges on imaging. (A) Meninges that remain uncut float up as shown by the white arrow. In this example, we see extensive protrusion of the meninges, because we did not remove them before agarose embedding. Usually, meninges remain uncut only for several difficult regions. (B) An example of bad slicing due to the presence of meninge between the corpus callosum and the upper part of the thalamus. In this case, bad slicing led to alternation of deep slice (329) and relatively normal slice (330). #78-329 stands for sample no. 78 section no. 329 in Brain/MINDS data portal (https://dataportal.brainminds.jp/marmoset-tracer-injection/viewer). (C) Another example of bad imaging. In a worse case, the pulvinar nucleus shown by the red arrow may entirely come off. (D) Another example of bad imaging. The meninge deep within the calcarine sulcus is difficult to remove. The shadows seen in sections 469 and 470 are caused by the floating meninges that came under the objective.

We try to manually extract them by fine forceps when we notice.

### 3. Auxiliary histological techniques

1. Section retrieval: The tissue slices can be later recovered to perform various histological staining. With training, it is possible to accurately align these sections in order of slicing. Trace the blood vessels across cortical layers for alignment (Fig. 3).
2. Cut off excess agarose from each section for better handling. This process needs not to be excessive. Actually, the agarose serves to hold tissue segments in place without interfering the histological processing.
3. [for Backlit imaging and Nissl staining] Mount the sections onto a slideglass and dry. Rehydrate the section with PBS and place the coverslip for imaging. With appropriate lighting, myelination pattern can be observed with no staining (Fig. 4). Remove the coverslip and proceed to Nissl staining. This way, you can obtain both histological data for the same tissue section (Fig. 4).
4. [for BDA imaging] The sections are first treated with methanol for permeabilization. Tyramide Signal Amplification (TSA) method is used to enhance the BDA signal to be fluorescently detected^17^. See Table 1 for detailed protocol.
5. [for smFP staining] The ‘spaghetti monster’ fluorescent proteins (smFPs) are a family of non-fluorescent GFP variants with multiple epitope tags^18^. They provide excellent fluorescent signals upon immunodetection and suitable as the companion tracers to the fluorescent tracers. See Table2 for our AAV constructs available from addgene.

**Fig. 3:**
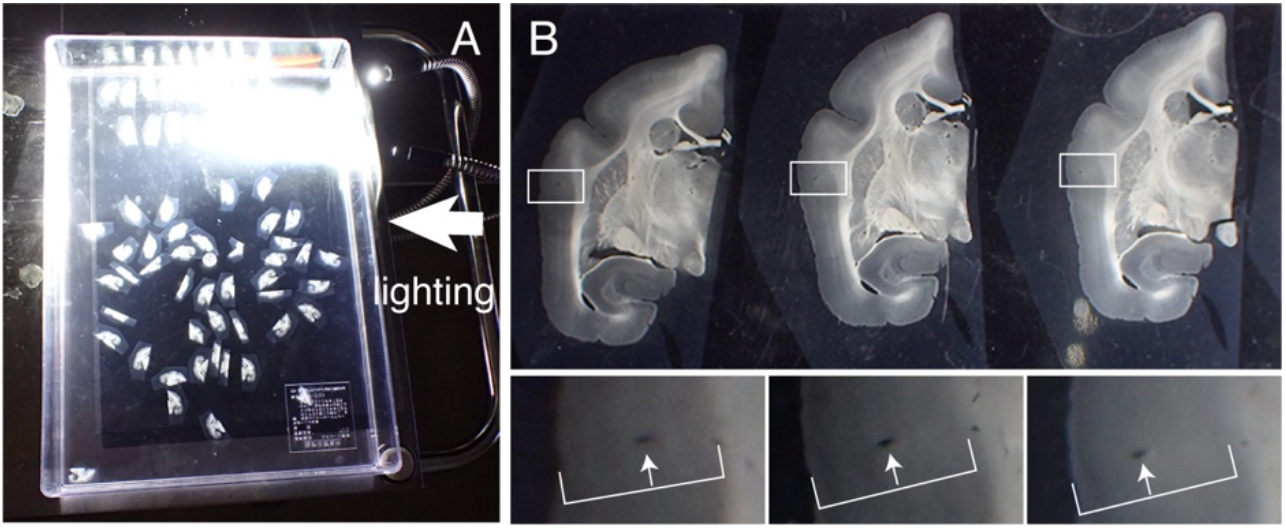
Alignment of tissue sections in order. (A) Up to 50 marmoset coronal sections can be correctly aligned in order in a plastic container (31×22.5cm). To visualize the detailed structures, the container is placed on a black paper and lighted from the side. These sections are first roughly aligned in order and then subject to precise alignment. (B) The precise alignment uses the blood vessel as the marker. The cerebral cortex contains many blood vessels that run vertically across cortical layers. They are identified as elongated holes that systematically change their positions within the cortical layers (white arrows). Using this method, even sections with 50 μm intervals can be accurately aligned. These blood vessel holes can be identified in the SPTP section images for confirmation.

**Fig. 4:**
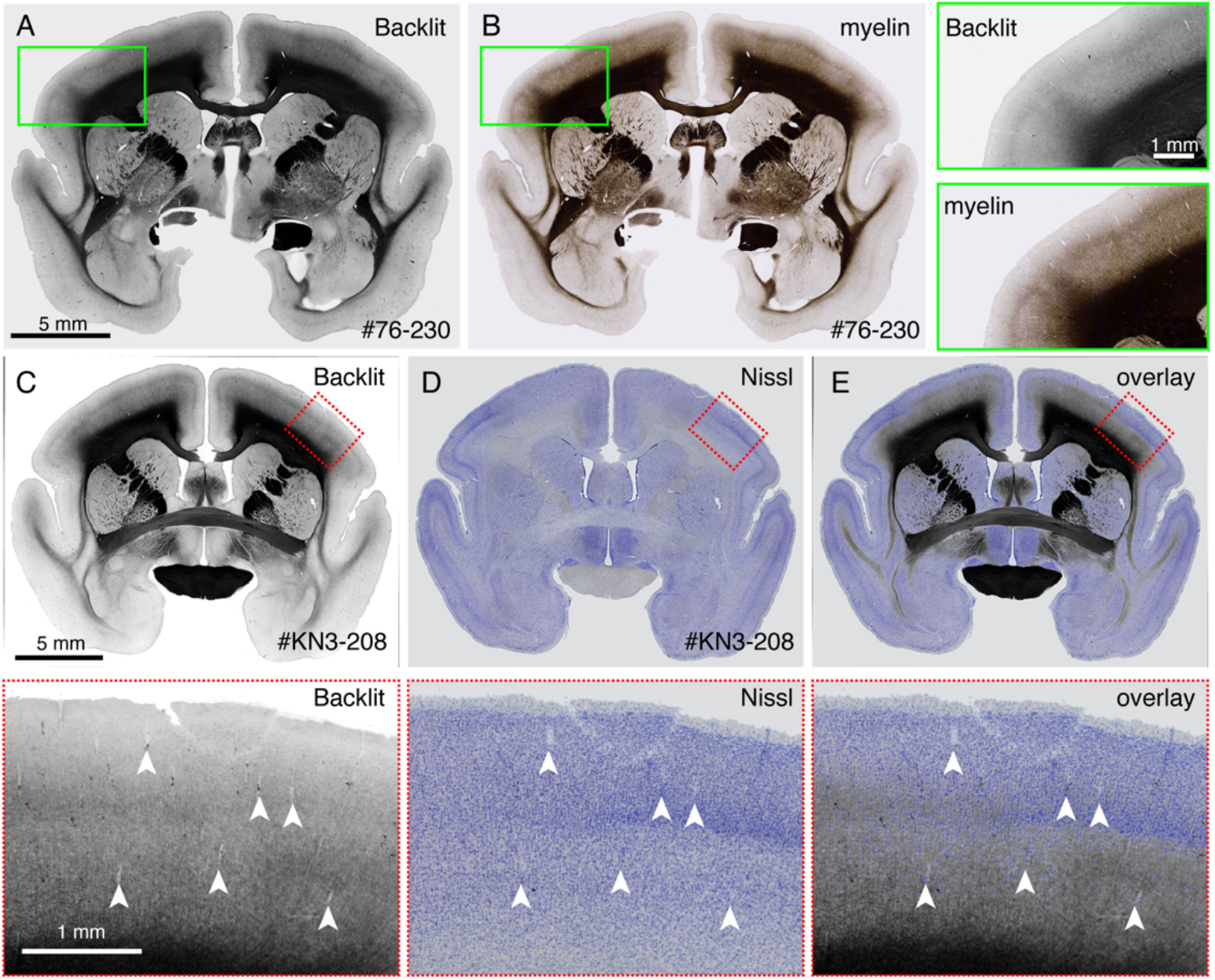
Backlit image can substitute for the myelin staining. (A and B) The identical section was used for backlit imaging and myelin staining. First, the section was mounted onto the slideglass, dried, rehydrated with PBS, coverslipped and imaged using Keyence microscope (BZ-X710). After removing the coverslip, the same section was used for myelin staining^27^ and imaged using the Keyence microscope. The backlit image was registrated to the myelin image using bUnwarpJ plugin of ImageJ. The green boxes show the magnified views of each image. Note that these images show almost identical patterns except that the myelin staining visualizes fibrous structures better. (C-E) The identical section was used for backlit imaging and Nissl staining. The low-threshold mask for the backlit image was first registrated to the low-threshold mask for the Nissl image using bUnwarpJ plugin and the original image was transformed using the same parameter. Note that the blood vessels (white arrowheads) are well matched between the two images. Because these two images are for the same section, the matching is almost perfect, and you can directly compare the myelin and Nissl patterns for identification of cortical layers.

**Table 1:** BDA fluorescent staining and anti-myc fluorescent staining protocol.

|  |  |
| --- | --- |
| Remove agarose |  |
| TBS wash | 10 min x 2 |
| 1% H<sub>2</sub>O<sub>2</sub> in Dent's solution | 10 min |
| TBS wash | Brief |
| 0.5% TNB blocking | 1hr |
| StAvHRP (1:4000) in TNB | 2 over night |
| TNT wash | 10 min x 3 |
| TSA Biotin(1:4000) in 0.1M borate(pH8.5) + 0.003%H<sub>2</sub>O<sub>2</sub> | 2hrs |
| TNT wash | 10 min x 3 |
| Cy3-streptavidin (1:1000) in TNT | 3hrs |
| TNT wash | 10 min x 2 |
| TBS wash | Keep the section until mounting |
| Mount section onto slideglass |  |

|  |  |
| --- | --- |
| Remove agarose |  |
| TBS wash | 10 min x 2 |
| blocking in IB | 1hr |
| Anti-Myc (1:4000) in IB | 2 over night |
| TNT wash | 10 min x 3 |
| Anti-mouse Cy3 (1:1000) in TNT | 3hrs |
| TNT wash | 10 min x 2 |
| TBS wash | Keep the section until mounting |
| Mount section onto slideglass |  |

**Table 2:**
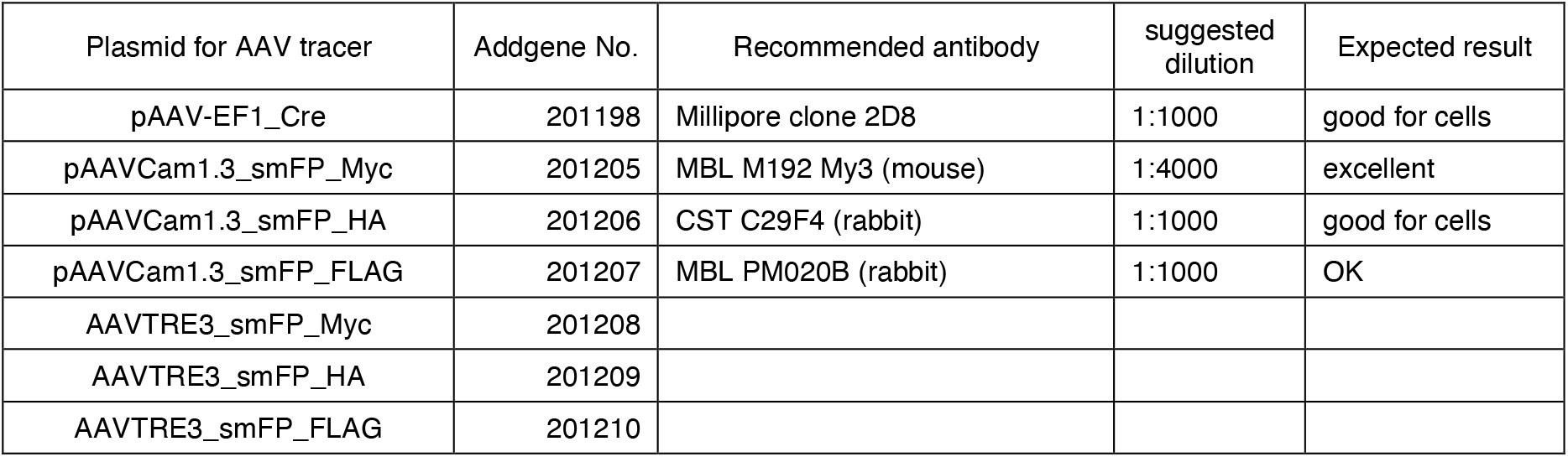
List of addgene plasmids available for production of non-fluorescent tracers. The cre construct targets the nucleus and suitable for retrograde tracing enveloped by AAV2 retro. smFP_HA construct is good for cell body detection (also for retrograde).

## REPRESENTATIVE RESULTS

In our typical set-up, a whole brain of an adult marmoset can be imaged at the resolution of ∼1.3×1.3 μm/pixel with 50 μm section interval in about one week. This amounts to ∼650 coronal images in three channels after image stitching. With 16x objective lens (Nikon 16xW CFI75

LWD; NA=0.80), the field of view of a single shot is about 1 × 1 mm. The image for the entire coronal surface is obtained by stitching these shots (Fig. 5A). The alignment in Z direction is excellent and a good 3D image is obtained by simply stacking the coronal image data (Fig. 5B and 5E). For standardization, STPT template for the marmoset brain^11^ can be used for 3D-3D registration (Fig. 5F). This data transformation is one of the key aspects of the “whole brain neuroanatomy”, in which a region of interest is assigned an absolute space coordinate, independent of anatomical annotation. Once the sample brain is registrated to the standard space, you can easily take out the subregions of interest for further analysis (Fig. 5F and 5G). In particular, the cortical regions can be transformed to a flatmap using a predetermined parameter (Fig. 5H).

**Fig. 5:**
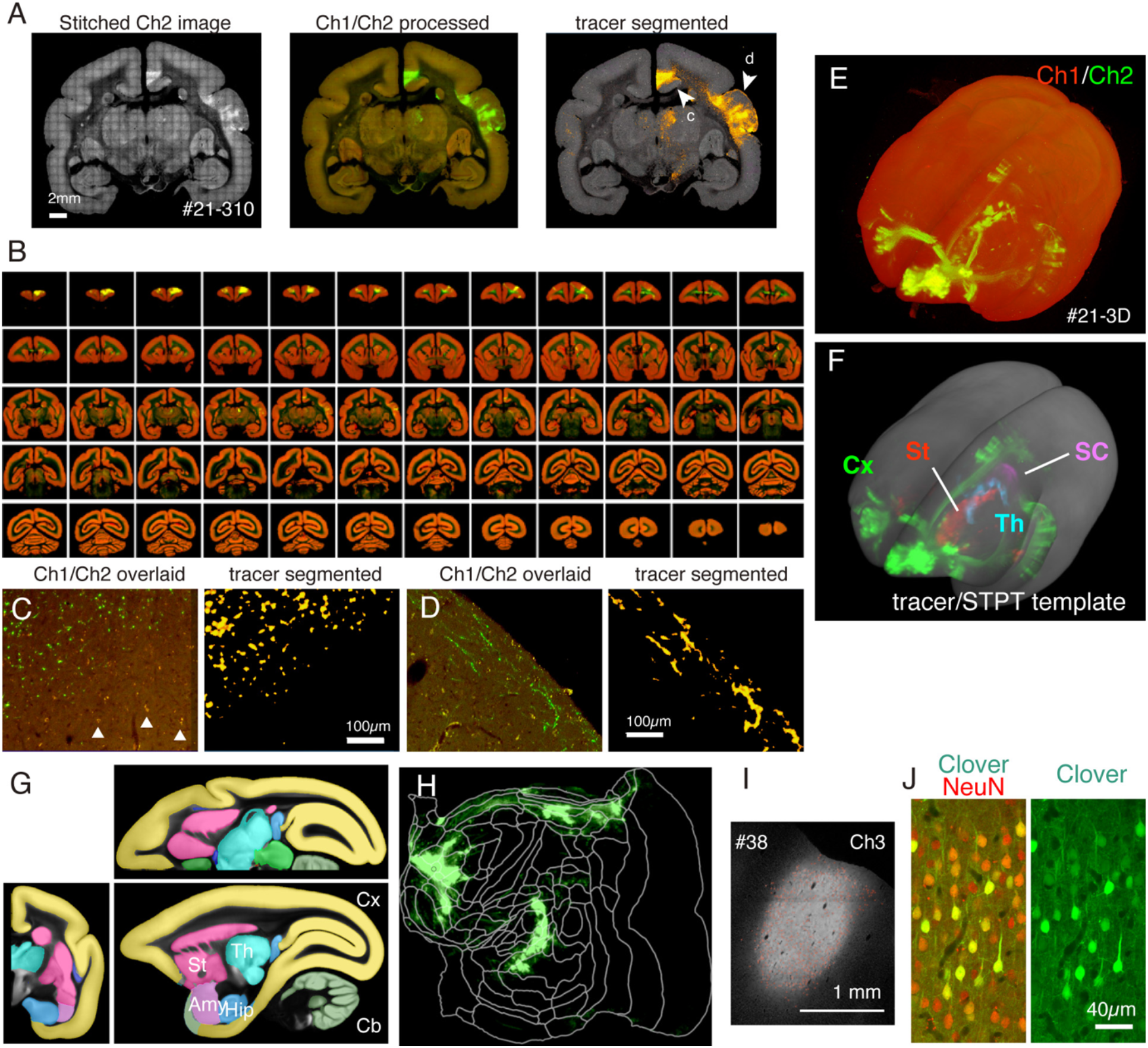
Typical result of STPT imaging. (A) A simple tiling of Ch2 (green) images of an example section (section 310 of sample #21). Without background correction, the borders for each tile are visible. In the middle column, the tiles were stitched with background correction for Ch1 (red) and Ch2 (green), respectively. To reduce the signals of lipofuscin (see panel C), Ch1 signals were subtracted from Ch2 and shown green. To reduce the lipofuscin signals of Ch1, “Remove Outilers..” command was used before stitching by ImageJ. In the right column, The tracer signals were segmented by image processing pipeline^11^. The arrowheads show the locations where the magnified views are shown in panels C and D. (B) Overview of serial sections for sample #21. STPT generated 635 high resolution coronal images for this sample. (C) A high magnification view shown by arrowhead c in panel A. This is a simple overlay of Ch1 (red) and Ch2 (green) with no further processing. The triangles show lipofuscin fluorescence signals, which show widespread spectrum. The tracer segmentation algorithm accurately distinguishes the tracer signals from the lipofuscin background despite very similar shape (right panel). (D) Another example of high magnification view. Note that fine axon fibers in layer 1 can be well visible. (E) The 3D reconstruction of original STPT images. The 635 low-resolution coronal images as shown in panel B were used as the tiff-stack for 3D visualization using fluorender^28^. (F) The 3D reconstruction of the segmented tracer signals registrated to the STPT template (grey). The tracer signals in different brain regions were shown by different colors. These regions were cut out by using the annotation shown in panel G. (G) STPT template overlaid with annotation of different brain regions. (H) Cortical tracer signal shown in panel F was shown in the form of flatmap. (I) Identification of the injection site by the Ch3 fluorescence, which is less sensitive to the tracer fluorescence and remain unsaturated. (J) Staining the section around the injection center with NeuN antibody showed that approximately 30 % of the neurons show strong expression of Clover green fluorescence. Cx; cortex, St; striatum, Th; thalamus, SC; superior colliculus. Amy; amygdala, Hip; hippocampus, Cb; cerebellum.

The sections generated during the imaging can be used for various histological purposes. As shown in Fig. 4A and 4B, the backlit imaging with no staining provides a very similar pattern to the authentic myelin staining. This can be a superb alternative to myelin staining. Furthermore, if the backlit image is obtained before Nissl staining, the same section can be used to obtain the patterns of both myelin and Nissl staining, thus, providing a useful information to identify areas and layers (Fig. 5C-E). These sections can also be used for immunological staining. In Fig. 5J, the section around the injection center was counterstained with NeuN antibody to estimate the transduction efficiency of the AAV virus. It is also a good strategy to inject non-fluorescent tracers in addition to fluorescent tracers and detect them histologically after section retrieval. In our previous study, we combined anterograde green tracers with retrograde “cre” vector, which was later detected by anti-cre antibody^10^. In an example of Fig. 6A-E, we injected BDA to the contralateral side of green tracer (clover) and fluorescently detected. Note that the red BDA signals can be registered to the TissueCyte image to be localized in the whole brain coordinate. In another example of Fig. 6F-I, smFP-myc was injected to the parietal area (Fig. 6G), while the green tracer was injected to the frontal area. This way, multiple tracers can be injected into the same animal without interfering the TissueCyte imaging. A big advantage of using the STPT sections for additional staining is that the relationship between the fluorescent and non-fluorescent tracers can be determined for the same brain. As such, we were able to determine the reciprocity of corticocortical projections at high-precision^10^. Another advantage is that the 3D coordinates of the stained sections can be mapped backed to the STPT data, and then to the standard template. Thus, it may not be needed to use all the retrieved sections for staining. We can select sections for staining to add further context to the STPT data for better interpretation.

**Fig. 6:**
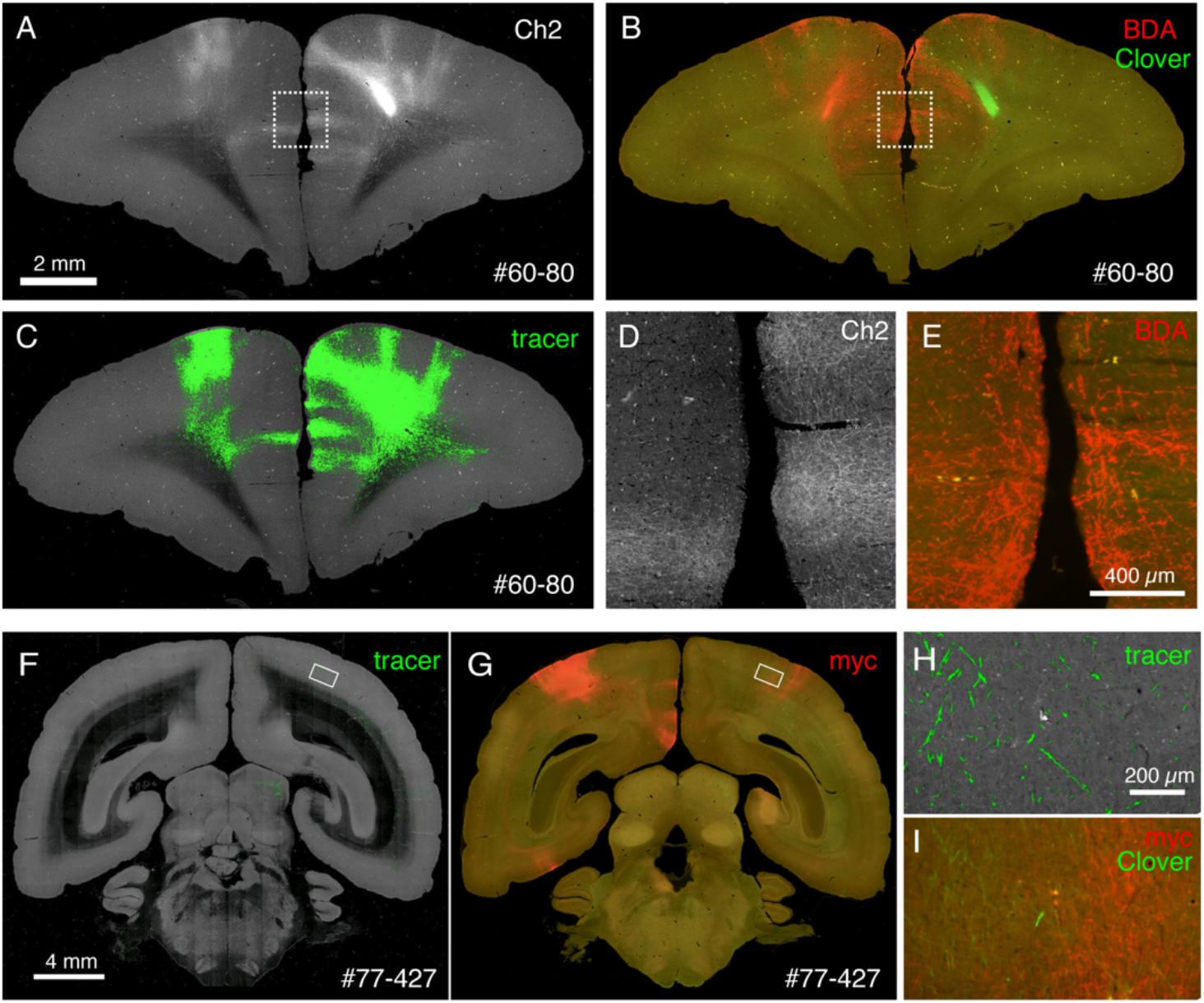
Post-STPT histology showing multiple non-fluorescent tracers. (A-C) Comparison of STPT image with BDA-stained image. In this sample, BDA is injected into the contralateral side of Clover injection. Panels A and C shows the Ch2 image and tracer segmentation of the STPT data. Panel B shows the post-STPT staining for BDA. Clover fluorescence is diminished due to methanol treatment of the section. (D and E) The dotted boxes in panels A and B are magnified. (F and G) Comparison of STPT image with anti-myc tag antibody staining. In this sample, Clover injection is to the PFC, whereas the AAV-smFP-myc is injected into the contralateral parietal cortex. The white rectangles are magnified in panels H and I. (H) Tracer segmentation is shown in green. (I) The myc staining is shown in red. The Clover fluorescence is shown in green. The green signals in panels H and I are present in a similar position but not identical, because STPT retrieves only ∼10 μm optical section.

## DISCUSSION

In this article, we explained the practical solutions to handle marmoset brains for whole brain processing as well as auxiliary histological techniques that enhance the utility of STPT technique. The strength of “whole brain neuroanatomy” using STPT is that you can obtain the 3D coordinates of any region of interest, whether it is anatomically annotated or not. By high precision 3D-to-3D registration, it is possible to transform these coordinates to a standard template for overlay of multiple datasets. This way, the standard template serves as the medium of data integration. This was an essential aspect of our PFC mapping project^10^, in which we analyzed data obtained from many different individuals. Furthermore, the information mapped to the standard template can be compared with various data that have already been mapped, whether it’s Nissl, myelin patterns, tracer data, MRI (including diffusion MRI) data or anatomical annotations^11^. Importantly, it can also be compared with future data that will be gained by yet-to-emerge technology. There currently exist multiple templates for the marmoset brain, which are based on Nissl staining^19^, MRI^19–24^ and STPT^11^. But they can be transformed to each other’s coordinates based on image contrasts and precalculated parameters^11^. We believe that data integration at whole brain scale across studies contribute to better understandings of the brain as a system. The prerequisite for this strategy to work is the reliable data acquisition across the brain. Below, we discuss potential problems associated with the current protocol.

[Technical considerations]

### > Imaging problems

One of the most vulnerable processes of STPT imaging is the tissue slicing. As we have mentioned above, the meninges often remain uncut and can interfere with the slicing. In particular, the pulvinar nucleus and the superior colliculus are two most affected brain regions: depending on the thoroughness of meninge removal and embedding in agarose, they can be stripped off from the tissue block during slicing. Another concern is flip back of sliced sections onto the tissue block, which sometimes occurs when the sections remain attached even after slicing. It can be minimized by shaping the block so that the blade cuts obliquely at the very end.

By imaging deep into the tissue block, STPT avoids the bumpiness of its surface. Whereas the fluorescent signals can pass through the cortical region rather easily, they are highly diminished in the myelinated regions. Therefore, the imaging depth needs to be carefully determined to balance between the consistent imaging across the entire block surface and the brightness of the signals in the myelin-rich region, such as the white matter. In our set-up, we usually aim at 25-35 μm from the surface. It also needs to be cautioned that the cut surface may become unevenly shrunk after long hours of storage. To minimize the imaging area, we split the imaging session into 20-30 runs with different stage settings for 5-6 days. We either make the interval between runs to be less than two hours or confirm the surface depth and adjust the stage height before each run.

### > On BDA staining

In our protocol, we amplify the BDA signal by TSA method^13^. This method is highly effective and can detect anterogradely transported BDA signals even at relatively low resolution (e.g., Fig. 6B). TSA biotin is commercially available from Akoya Biosciences, but the home-made solution shows much better enhancement. On the other hand, the dilution of the antibody and solution requires careful adjustment to obtain the optimal result. Pretreatment of the section by methanol solution is critical. Without pretreatment, the BDA signals is barely detected in the myelinated axons.

### > Identification of the injection site

When we use an anterograde tracer, it is often difficult to identify the exact site of injection, because of saturation of the fluorescence signals. In our TissueCyte set-up, we use the blue channel to identify the infected cells (Fig. 5I). Even when the red and green channels are saturated, each individual infected neurons are usually detectable in the blue channel. This is also true with the Allen Mouse Brain Connectivity Atlas^1^. Examination of the cells of origin is important because the infection sometimes involves only particular layers. We encountered such partial infections rather frequently for the Mouse Brain Connectivity Atlas, perhaps because of utilization of iontophoresis method^25^. The lateral spread of the viral tracers could be more or less variable depending on the injections. This variability can potentially affect the result of tracing and needs careful normalization.

### > On registration

The successful registration of the obtained 3D image to the standard template is a key process of the whole brain neuroanatomy. Registration of the STPT image to the STPT template can be pretty accurate and we observed only deviances of a few voxels (50μm isocubic) for the borders with a high image contrast^10^. Still, there is a limit to what registration can do. Because of the meninge removal process, the STPT samples generally have gaps between the hemispheres and between the cortex and midbrain/hindbrain, whereas brain tissues are tightly packed in the in vivo MRI images. Such differences are difficult to adjust by registration. Marmoset cortex is mostly devoid of sulci, but the intraparietal sulcus is very deep in some Individuals. Such a sulci will be lost (top-to-top fusion occurs) upon registration. Although registration is a powerful technique, it is necessary to go back to the raw data for confirmation of the obtained result.

## ACKNOWLEDGMENTS

We deeply thank technical staffs in Yamamori lab and animal facilities (RRD) for help. We thank the RIKEN CBS-Olympus Collaboration Center for the technical assistance with confocal image acquisition. This work was supported by the program for Scientific Research on Innovative Areas (grant number, 22123009) from MEXT, Japan, by the program for Brain Mapping by Integrated Neuro technologies for Disease Studies (Brain/MINDS: JP15dm0207001 to T.Y.) from AMED, Japan, and by JSPS KAKENHI Grant Number 24K09678 to A.W.

